# A new pipeline for the normalization and pooling of metabolomics data

**DOI:** 10.1101/2021.07.16.452593

**Authors:** Vivian Viallon, Mathilde His, Sabina Rinaldi, Marie Breeur, Audrey Gicquiau, Bertrand Hemon, Kim Overvad, Anne Tjønneland, Agnetha Linn Rostgaard-Hansen, Joseph A. Rothwell, Lucie Lecuyer, Gianluca Severi, Rudolf Kaaks, Theron Johnson, Matthias B. Schulze, Domenico Palli, Claudia Agnoli, Salvatore Panico, Rosario Tumino, Fulvio Ricceri, Monique Verschuren, Peter Engelfriet, Charlotte Onland, Roel Vermeulen, Therese Haugdahl Nøst, Ilona Urbarova, Raul Zamora-Ros, Miguel Rodriguez-Barranco, Pilar Amiano, José Maria Huerta, Eva Ardanaz, Olle Melander, Filip Ottoson, Linda Vidman, Matilda Rentoft, Julie A Schmidt, Ruth C Travis, Elisabete Weiderpass, Mattias Johansson, Laure Dossus, Mazda Jenab, Marc J Gunter, Lorenzo Bermejo, Dominique Scherer, Reza M Salek, Pekka Keski-Rahkonen, Pietro Ferrari

## Abstract

Pooling metabolomics data across studies is often desirable to increase the statistical power of the analysis. However, this can raise methodological challenges as several preanalytical and analytical factors could introduce differences in measured concentrations and variability between datasets. Specifically, different studies may use variable sample types (e.g., serum versus plasma) collected, treated and stored according to different protocols, and assayed in different laboratories using different instruments. To address these issues, a new pipeline was developed to normalize and pool metabolomics data through a set of sequential steps: (i) exclusions of the least informative observations and metabolites and removal of outliers; imputation of missing data; (ii) identification of the main sources of variability through PC-PR2 analysis; (iii) application of linear mixed models to remove unwanted variability, including samples’ originating study and batch, and preserve biological variations while accounting for potential differences in the residual variances across studies. This pipeline was applied to targeted metabolomics data acquired using Biocrates AbsoluteIDQ kits in eight case-control studies nested within the European Prospective Investigation into Cancer and Nutrition (EPIC) cohort. Comprehensive examination of metabolomics measurements indicated that the pipeline improved the comparability of data across the studies. Our pipeline can be adapted to normalize other molecular data, including biomarkers as well as proteomics data, and could be used for pooling molecular datasets, for example in international consortia, to limit biases introduced by inter-study variability. This versatility of the pipeline makes our work of potential interest to molecular epidemiologists.

## 1. Introduction

Metabolomics is a powerful tool for investigating candidate etiological pathways for chronic diseases^[1–4]^. Using either untargeted or targeted (via sets of pre-defined annotated metabolites) approaches, prior metabolomics studies have identified metabolites associated with the risk of several chronic conditions, including type-2 diabetes (T2D)^[5]^, cardiovascular diseases (CVD)^[6]^, and cancer^[7–9]^. Metabolomics has also been used to characterize specific signatures of anthropometric measures and lifestyle exposures, including body mass index (BMI)^[7,10]^, adherence to a Mediterranean diet^[6]^ and coffee consumption^[5]^, as a way to investigate candidate biological mechanisms underpinning the relationship between these exposures and chronic diseases.

Like for other -omics technologies, pre-processing of metabolomics data is critical before relating them to phenotypes, such as cancer endpoints or lifestyle exposures^[11,12]^. After a matrix of *p* metabolites (or features) measured in *n* samples has been generated, pre-processing usually involves (i) feature and sample filtering, where low-quality features and samples are excluded (ii) data imputation, to take care of missing values and (iii) data normalization, to correct for sources of unwanted variation in metabolomics data, such as batch effects and other factors related to the handling of samples^[11,13–16]^. Following the success of data acquisition efforts in large-scale epidemiological investigation, collaborative consortia have been put in place, offering the possibility to pool metabolomics data acquired in different studies in order to increase sample size and range of biological variation, and eventually enhance statistical power of the analysis. However, pooling metabolomics data across studies raises methodological challenges as several preanalytical and analytical factors can induce differences in metabolite measurements and unwanted variability between datasets. Specifically, sample types (e.g., serum versus plasma), fasting status of the participant, and any other elements related to sampling conditions, samples’ treatment and storage represent preanalytical factors, while analytical factors include information on the organization of samples in batches, the acquisition instrument, the acquisition time (i.e., time at which the sample was assayed) and the laboratory^[17]^. Correcting for these sources of variations is crucial in order to conduct accurate statistical analyses on pooled datasets.

Data on common quality controls assayed in all studies and/or reference assay data from a subset of samples in each study can be used for normalization^[17,18]^. However, these data are not always available in large international investigations and consortia. Accordingly, we developed a pipeline for the normalization and pooling of metabolomics data acquired in different studies that does not require data on quality controls or reference assay data, which covers four main steps. First, data cleaning identified and removed features and samples exceeding certain thresholds of missingness and outlying samples^[11,16]^. Second, the remaining missing values were imputed within each study using information on limits of detection and quantification when available and appropriate, and measurements were log-transformed to reduce skewness. Third, the principal component partial R-square (PC-PR2) technique was implemented to identify sources of variation in the metabolomics data^[13]^. Last, mixed effect models were used to correct for unwanted variability while preserving biological variability^[14]^. The ComBat method^[19]^ implemented in the R sva package^[20]^ was also implemented for sake of comparison. Our pipeline was applied to targeted metabolomics data acquired in eight case-control studies nested within the European Prospective Investigation into Cancer and Nutrition (EPIC)^[21]^. Comprehensive analytical and graphical examinations of measurements were performed to assess whether different normalization approaches improved the comparability of metabolomics data. For illustration, metabolomics data were pooled and related to study participants’ BMI.

## 2. Results

### 2.1. Description of the study population

Targeted metabolomics data acquired within the EPIC study and centralized at the International Agency for Research on Cancer (IARC) included 16,060 pre-diagnostic blood samples originating from eight case-control studies nested within EPIC (details in Section 4.1) on seven types of cancer: breast cancer (one study denoted by BREA; n=3,172 samples)^[8]^, endometrial cancer (ENDO; n=1,706)^[22]^, gallbladder cancer (GLBD; n=112), liver cancer (LIVE; n=596)^[24]^, kidney cancer (KIDN; n=1,213)^[23]^, prostate cancer (PROS; n=6,020)^[9,25]^, and colorectal cancer (two studies denoted by CLRT1 and CLRT2; n=946 and n=2,295, respectively). As displayed in Table 1, samples collected at recruitment were assayed at IARC for BREA, LIVE, KIDN, PROS, and CLRT1, at the Helmholtz Zentrum (München, Germany) for CLRT2 and GLBD, and at the Imperial College London (UK) for ENDO. Across all studies, measurements of a total of 171 metabolites were acquired using either the AbsoluteIDQ p180 or the AbsoluteIDQ p150 (for CLRT2 only) commercial kit (Biocrates Life Science AG, Innsbruck Austria), following the procedure recommended by the vendor. As displayed in Table 1, samples were assayed on different Liquid Chromatography (LC) and Mass Spectrometry (MS) instruments across the different studies, but each study used one single pair of LC-MS instruments for all samples. Samples were mostly either serum or citrate plasma, and samples within one study all originated from the same type of blood matrix, except in BREA and GLBD where samples from Swedish participants originated from a different blood matrix compared to the other participants. For these two studies, samples assayed within each batch all originated from the same blood matrix (not shown). Samples were assayed between 2013 and 2018. The pipeline detailed in Section 4.2 was applied to the (n×p) matrix with n=16,060 samples and the p=118 metabolites measured in all studies. Specifically, they included 13 metabolites (amino acids) measured by a quantitative LC*-*MS/MS method (Liquid Chomatography coupled with tandem massspectrometry) and 105 metabolites (76 glycerophospholipids, 12 sphingolipids, 16 acylcarnitines, and 1 hexose, the sum of six-carbon sugars) acquired by a semi-quantitative FIA-MS/MS method (Flow-Injection Analysis coupled with tandem mass-spectrometry, one-point calibration, no individual internal standards).

**Table 1.** Main characteristics of the study population.

| Acronym | Number of Samples | Matrix | Laboratory | MS Instrument | LC Instrument | Kit Used |
| --- | --- | --- | --- | --- | --- | --- |
| BREA | 3,172 | Citrate plasma<sup>1</sup> | IARC | SCIEX QTRAP 5500 | Agilent 1290 | p180 |
| CLRT1 | 946 | Citrate plasma | IARC | SCIEX Triple Quad 4500 | Agilent 1290 | p180 |
| CLRT2 | 2,295 | Serum | HZM<sup>3</sup> | SCIEX API 4000 | Agilent 1200 | p150 |
| ENDO | 1,706 | Citrate plasma | ICL<sup>4</sup> | SCIEX API 4000 | Agilent 1290 | p180 |
| GLBD | 112 | Serum<sup>2</sup> | HZM<sup>3</sup> | SCIEX API 4000 | Agilent 1200 | p180 |
| LIVE | 662 | Serum | IARC | SCIEX QTRAP 5500 | Agilent 1290 | p180 |
| KIDN | 1,213 | Citrate plasma | IARC | SCIEX QTRAP 5500 | Agilent 1290 | p180 |
| PROS | 6,020 | Citrate plasma | IARC | SCIEX Triple Quad 4500 | Agilent 1290 | p180 |

<sup>1</sup>except Swedish participants (n=101; EDTA plasma). <sup>2</sup>except for Swedish participants (n=14, heparin plasma). <sup>3</sup>Helmhotz Zentrum München. <sup>4</sup>Imperial College London

### 2.2. Data cleaning and imputation

For the exclusion of metabolites and samples exceeding a given threshold of missingness, we applied the method described in Section 4.2.1 with a threshold set to 20%, and with missing values defined as “fully missing” values only, i.e., not including out of measurable range values. Among the 118 metabolites originally retained for the analysis, the acylcarnitine C4-OH (C3-DC) was the only one with a fully missing value in more than 20% of the samples of at least one study (PROS), and was excluded. Among the 16,060 samples originally retained for the analysis, none was excluded because of exceeding 20% of fully missing values, eight were excluded because they were measured in batches with less than 10 samples, and two were excluded because they were considered as outliers after a principal component analysis (PCA). Thus, the final study population included 16,050 samples, for which measurements of 117 metabolites were included (Supplementary Table 1). Out of the 1,877,850 corresponding measurements, 1,066 were fully missing and 63,564 were out of the measurable range: specifically, 63,044 were below a known LOD (limit of detection, applicable to acylcarnitines, glycerophospholipids, hexose and sphingolipids), 517 below a known LLOQ (lower limit of quantification, applicable to amino acids), 2 above a known ULOQ (upper limit of quantification) and 1 below an unknown LOD. All these 1,066 + 63,564 = 64,630 missing values were imputed as described in Section 4.2.2, and concentration values were log-transformed.

### 2.3. Identification of major sources of variations

As displayed in Figure 1 (left panel), the projection of the measurements on the first two principal components of the PCA were strongly clustered by study, suggesting the presence of systematic sources of heterogeneity across studies. The PC-PR2 method was applied to assess the proportion of the overall variation in the metabolomics data that was explained by a predefined list of variables, including (i) participants’ characteristics, i.e., study center, gender, case-control indicator, age, BMI, alcohol intake, smoking status, and (ii) three variables describing possible preanalytical and analytical sources of unwanted variations: fasting status, time of the day of blood collection, study and batch, with batch nested within study. As shown in Figure 2 (top panel), the PC-PR2 analysis indicated that these variables together explained more than 55% of the total variation of the metabolomics measurements before normalization. Study explained 31% of the total variation, while batch within study explained about 8%. Study center explained about 9% of the total variation, and gender, BMI and alcohol intake explained about 2%, 2% and 1%, respectively. Fasting status, time of blood collection, age at recruitment, smoking status and case-control status all explained less than 1% of the total variation.

**Figure 1.**
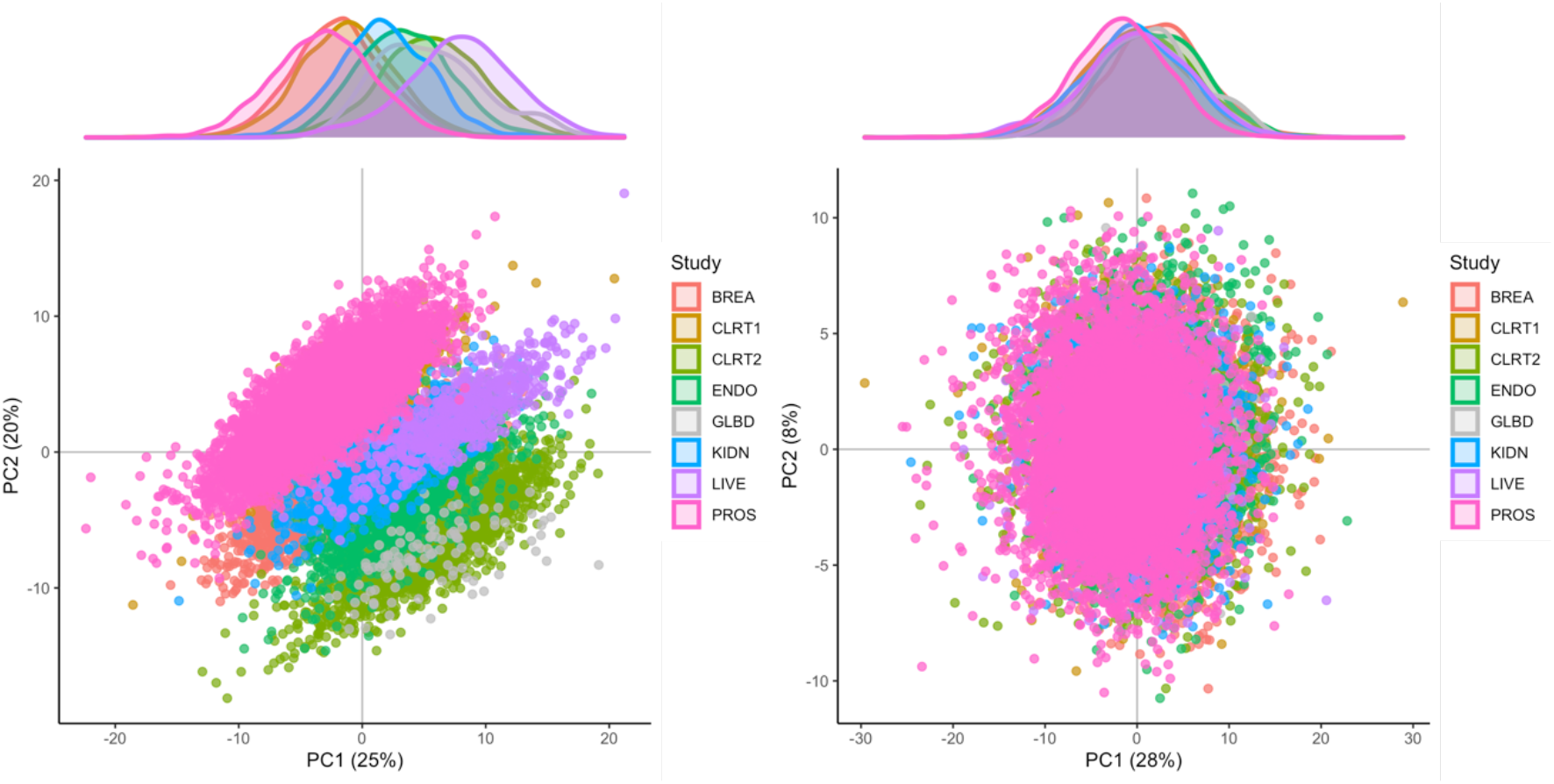
Results from the principal component analysis (PCA); left panel: PCA of the imputed data (i.e., before the normalization step); right panel: PCA of the normalized data.

**Figure 2.**
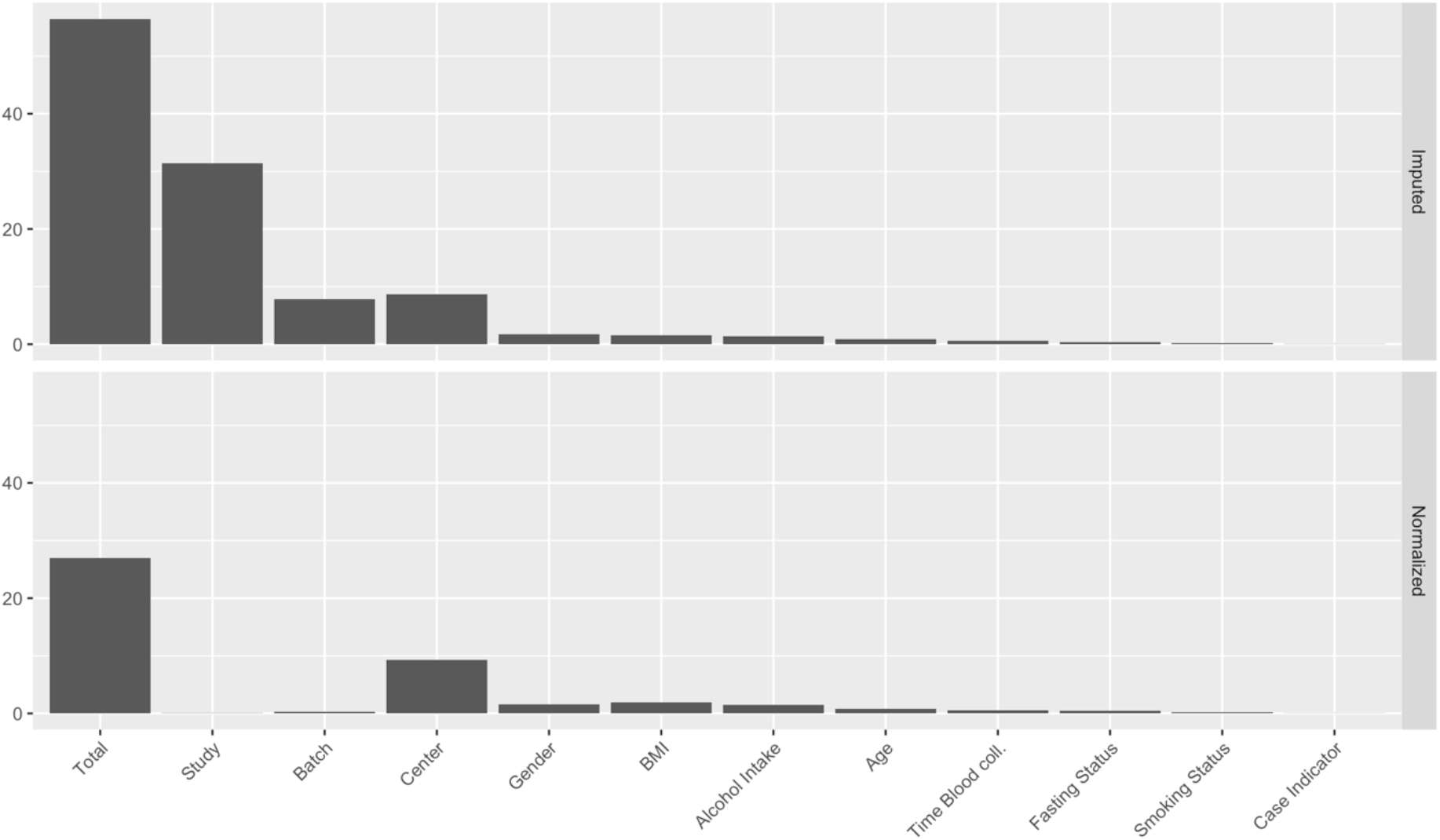
Results from the PCPR2 analysis of the imputed data (i.e., before the normalization step; top) and the normalized data (bottom).

### 2.4. Normalization of the measurements

Based on PC-PR2 analysis, metabolite concentrations were normalized using the method described in Section 4.2.4 to correct for variation due to study and batch, and preserve the variation due to study center, gender, BMI and alcohol intake. These latter four variables were all unequally distributed across studies and batches (not shown). They were included as fixed effects in matrix Z (Equation (2) in Section 4.2.4; otherwise some of the variation they explain would be removed because of the adjustment for study and batch), while study and batch within study were modeled as random effects in matrix X. Other variables studied in the PC-PR2 analysis were not included in matrix X or Z as they contributed very little to the total variation. Heteroscedastic metabolite-specific mixed models with a study-specific variance component were used, although homoscedastic models produced very similar results (not shown). The PCA of normalized data (Figure 1; right panel) indicated that the projections on the first two principal components were not clustered by study anymore, and measurements’ distribution largely overlapped. Data from PROS (men only) were slightly shifted to the left, and data from BREA and ENDO (women only) were shifted to the right, suggesting that the normalization preserved some variation due to gender overall. For illustration, the distribution of one semi-quantified metabolite, SM OH C22:1, was computed within batches and across studies, for the imputed and the normalized measurements (Figure 3). Imputed data displayed a study effect, with concentration levels of SM OH C22:1 in the CLRT2, ENDO, GLBD, KIDN and LIVE studies substantially larger than those in BREA, CLRT1 and PROS. A remarkable batch effect was observed within some studies, e.g., BREA. After normalization, the distributions were very similar across batches and studies. Again, the distribution was slighty shifted downward for concentration levels in PROS (men only), and upward in BREA and ENDO (women only), compared to the other five studies CLRT1, CLRT2, GLBD, KIDN and LIVE (which included both men and women), confirming that the normalization preserved some variation due to gender for this particular metabolite. The PC-PR2 analysis of normalized data (Figure 2, bottom panel) confirmed that normalization removed unwanted sources of variation (batch and study), but kept most variability attributed to participants’ characteristics. Complementary PC-PR2 analysis showed that blood matrix and LC-MS instruments contributed to less than 0.1% of the total variation after normalization (results not shown). Compared to our approach, ComBa^*[19]*^ produced very similar results for all metabolites with the exception of most acylcarnitines and the glycerophospholipid PC aa C40:1 (Supplementary Figure 1).

**Figure 3.**
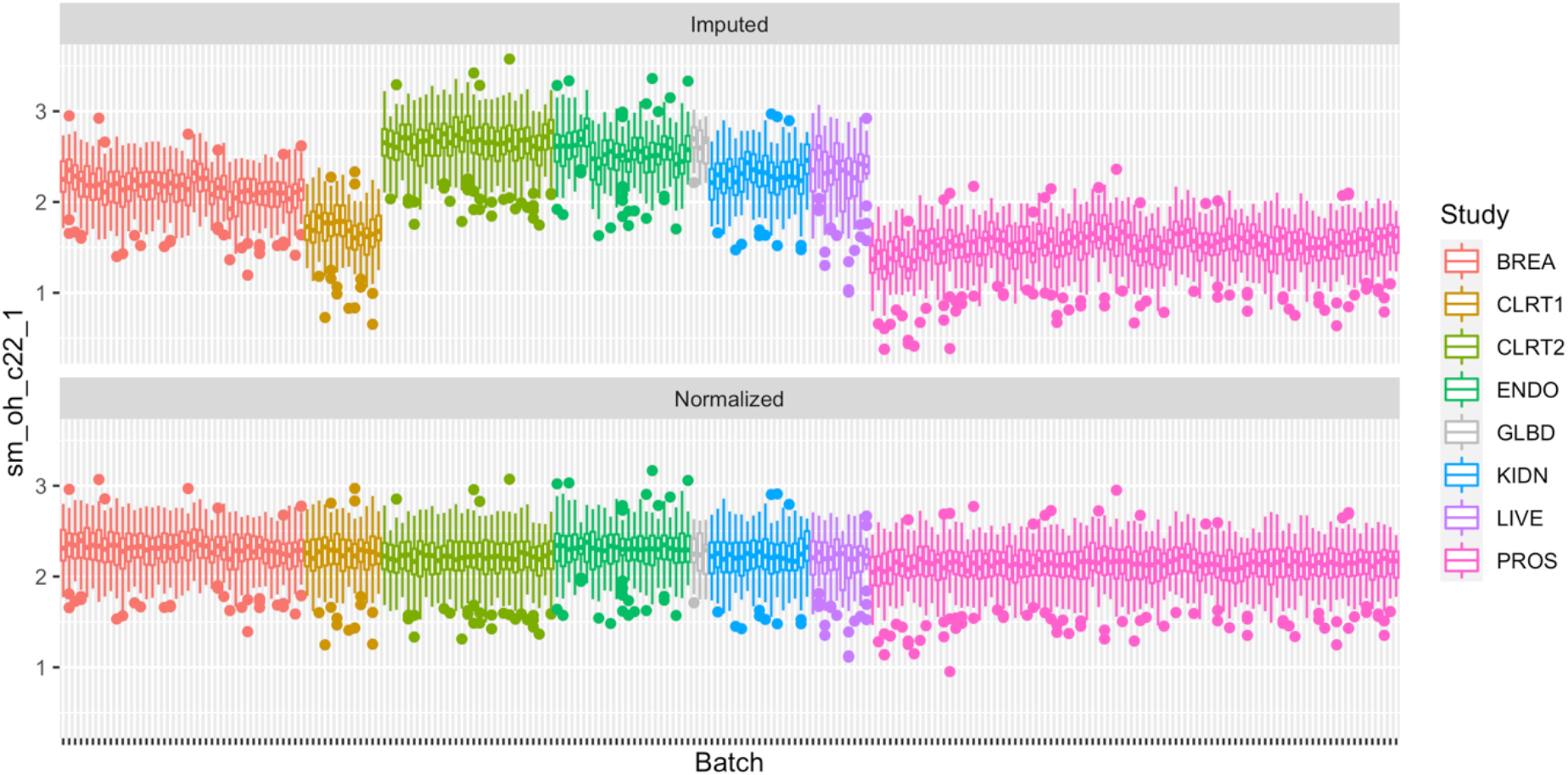
Boxplots of SM OH C22:1 within each of the eight case-control studies for the imputed data and the normalized data. Dots indicate measurements out of the interval [q1 - 1.5*IQR, q3 + 1.5*IQR] with q1 and q3 the first and third quartile, respectively, and IQR=q3-q1 the interquartile range.

### 2.5. Technical reproducibility of measurements before and after the normalization

Intra-class correlation (ICC) coefficients were computed for each metabolite to assess their technical reproducibility, using measurements from 2*219=438 duplicate samples, i.e., samples measured once in two different studies (2*147 samples; see Supplementary Table 2) or in two different batches of the prostate study (2*72 samples), as detailed in Section 4.3. Figure 4 shows the distributions of ICCs, for the semi-quantified (lipids, acylcarnitines and hexose) and quantified (amino-acids) metabolites, before and after normalization. The normalization shifted the distribution of ICCs upward for semi-quantified metabolites. The distribution of quantified metabolites did not shift as much, but the variability narrowed down, with no ICC value lower than 0.50. Figure 5 shows the effect of the normalization on the ICC of each individual metabolite (top), and on the average ICC for each class of metabolites (bottom). Before normalisation, 101 (86%) metabolites (92 semi-quantified, 9 quantified) had ICC values lower than 0.75, among which 38 (32%: 35 semi-quantified, 3 quantified) had ICC values lower than 0.5. After normalization, only twelve metabolites (10%: 9 semi-quantified, 3 quantified) had ICC values lower than 0.75, among which only two semi-quantified metabolites had ICC lower than 0.50. Also, class-specific averaged ICC values were consistently improved by the normalization, in particular for glycerophospholipids and sphyngomyelins. Similar results were observed when normalization was performed with ComBa^*[19]*^, yet ICCs were larger when using our approach, especially for acylcarnitines (Supplementary Figure 2). The same analysis was restricted to the 2*57=114 duplicate samples aquired in two studies from serum and citrate plasma, respectively. As displayed in Supplementary Figure 3, ICC values were lower than 0.5 for 69 metabolites (59%) and 4 metabolites (3%), before and after normalization respectively, with ICC values greater than 0.75 for 91 metabolites (78%) after normalization.

**Figure 4.**
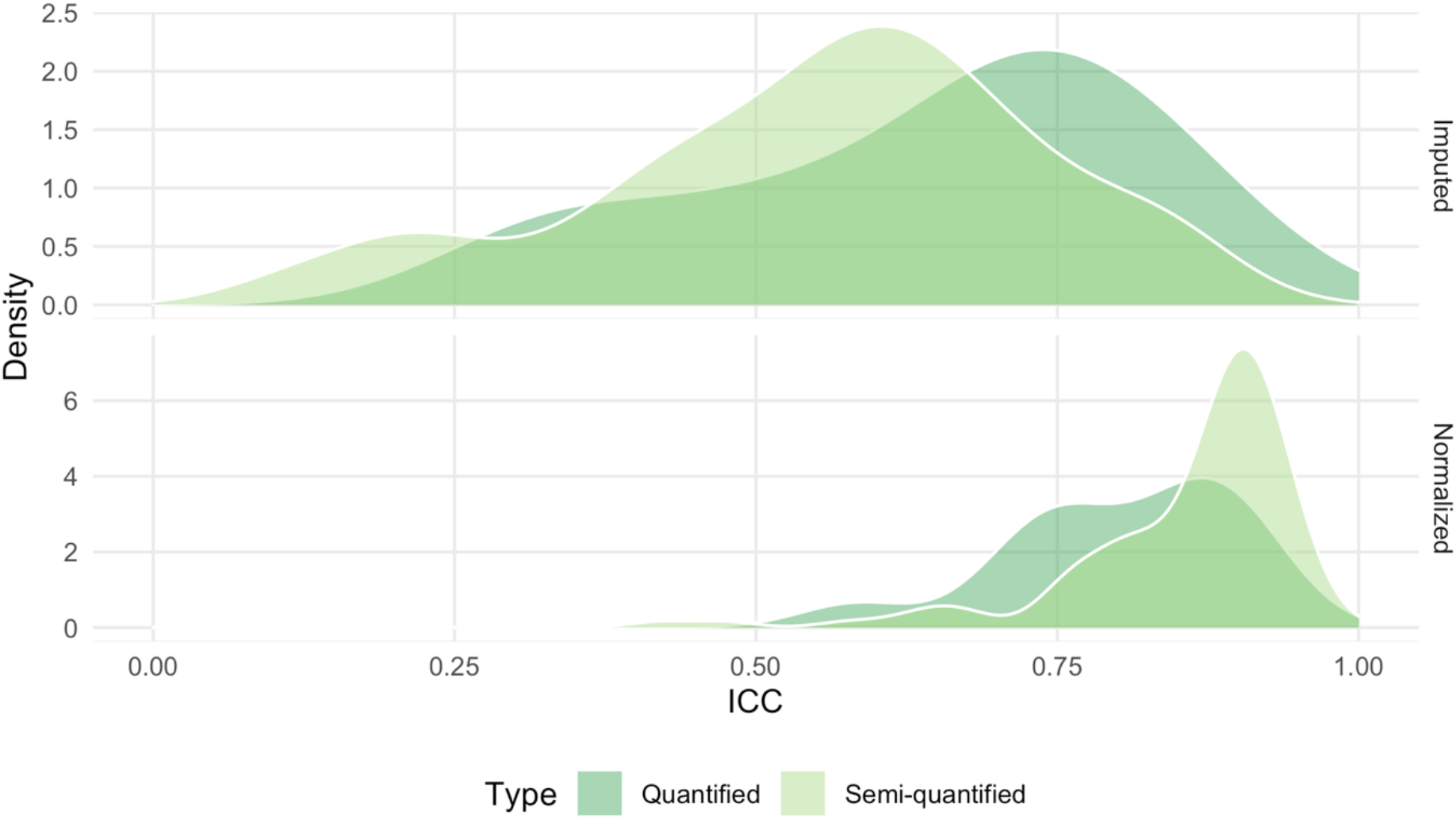
Distribution of the Intra-Class Correlation (ICC) coefficient for quantified and semi-quantified, before (top) and after (bottom) normalization.

**Figure 5.**
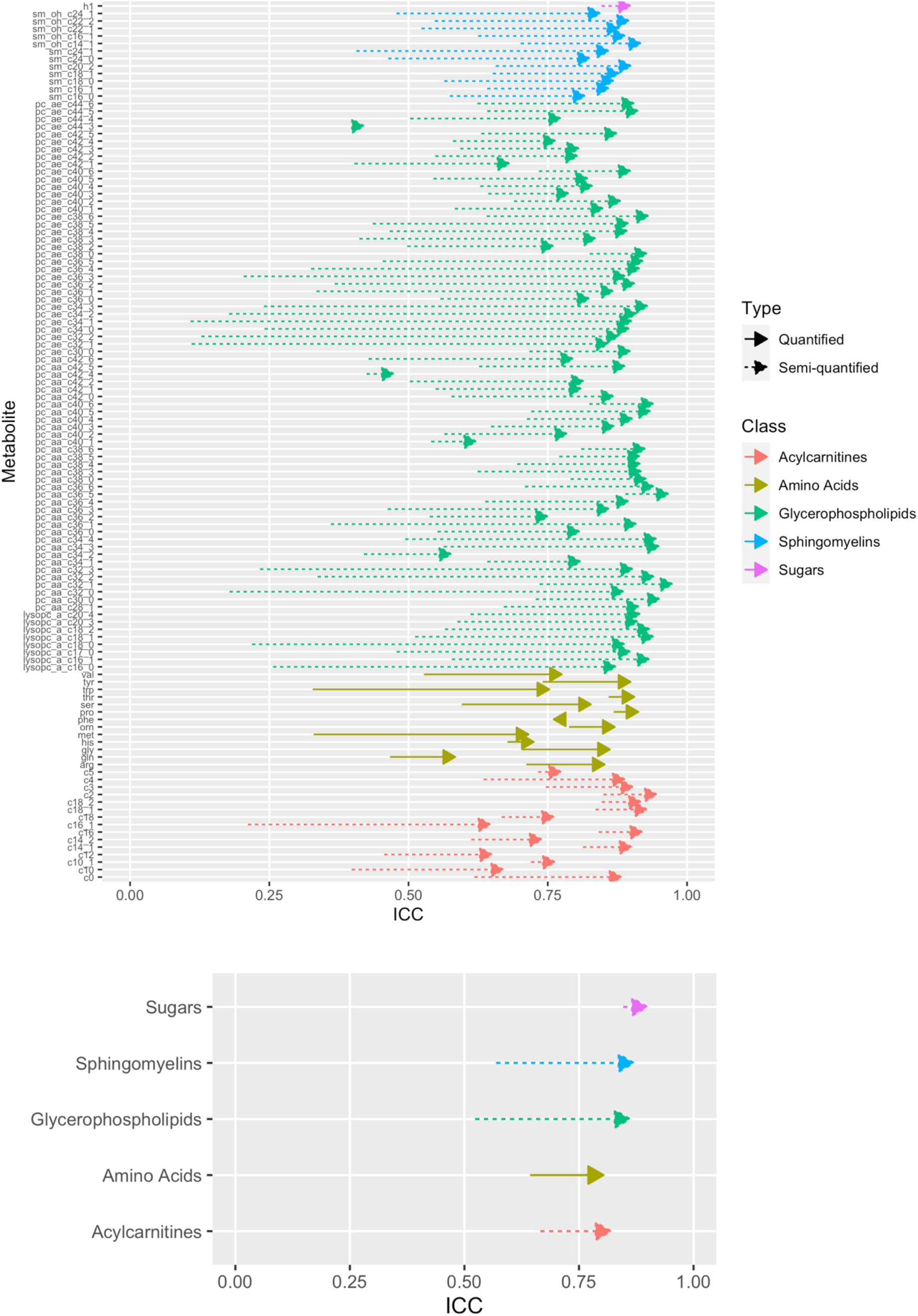
Metabolite-specific ICC values before and after normalization (top) and average ICC values for each class of metabolites before and after normalization (bottom). For each arrow, its origin represents the ICC value before normalization, and its peak represents the ICC value after normalization.

### 2.6. Impact of the normalization when relating a given phenotype to the metabolites

The relationship between the metabolites and BMI was assessed. The analysis was restricted to control samples to reduce collider bias, and one sample was randomly chosen among duplicates. For each of the 117 metabolites, Pearson correlation coefficients were computed between BMI and, in turn, the imputed measurements, the normalized measurements, as well as the normalized measurements produced by a simpler normalization approach, which corrected for study and batch effects without attempting to preserve variation due to study center, BMI, gender and alcohol intake. As displayed in Figure 6, most correlation values were above the line y=x, especially for values greater than 0.1: associations with BMI were stronger when using normalized data implementing our approach, compared to those observed with both non-normalized data and normalized data implementing a simple, yet incomplete, normalization approach.

**Figure 6.**
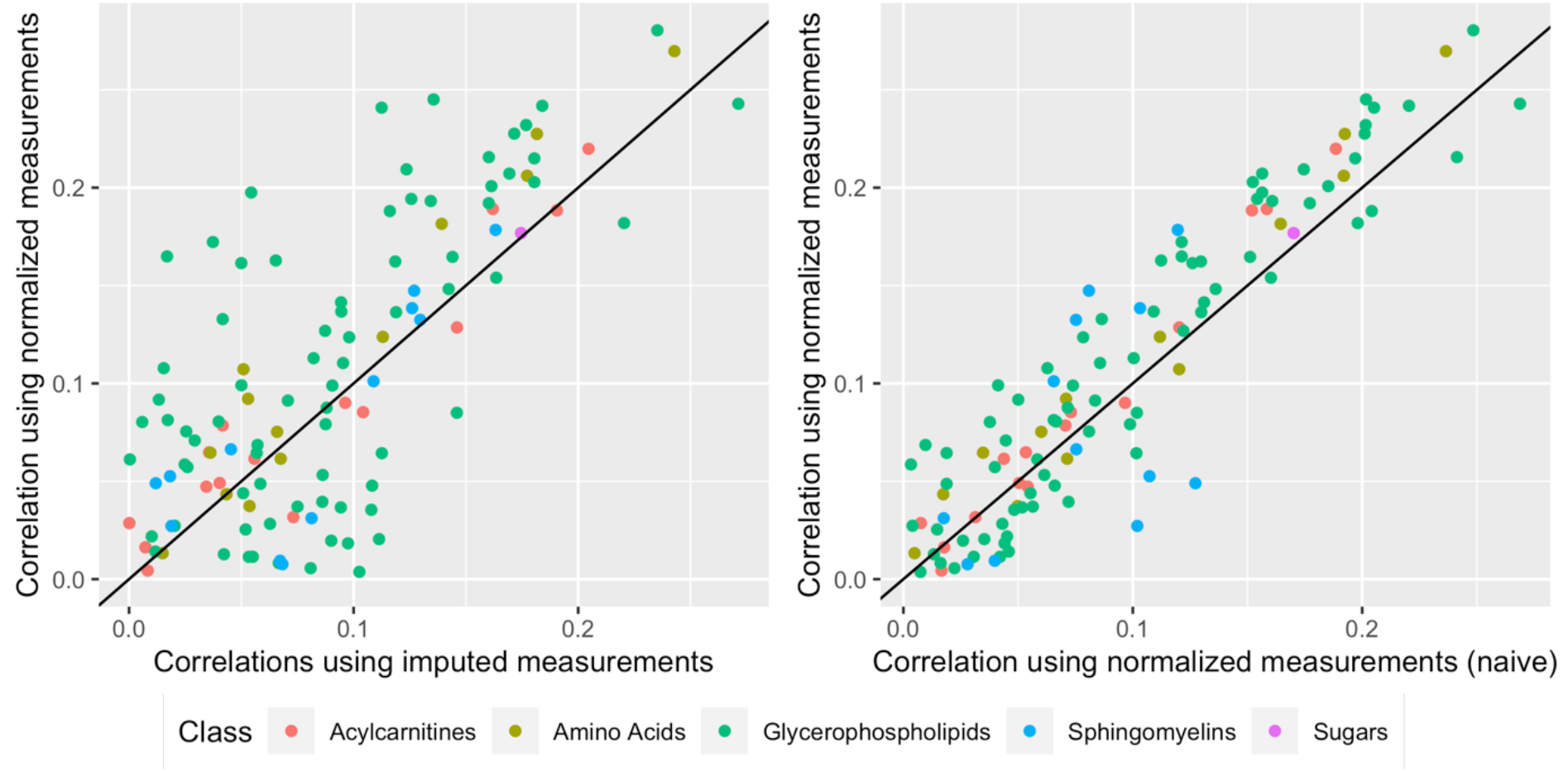
Correlations (absolute values) between BMI and the 117 metabolites in control samples. The *y*-axis represents values computed with normalized measurements produced by our approach, while the *x*-axis represents values computed with imputed (non-normalized) measurements (left), and normalized measurements produced by the simpler normalization approach (right), which corrected for study and batch effects without specifically attempting to preserve variation due to study center, BMI, gender and alcohol intake.

## 3. Discussion

In this work, a pipeline for the normalization of metabolomics data acquired in different studies was described. After a screening of informative metabolites and samples, the PC-PR2 method was used to identify major sources of variation in metabolomics data, and linear mixed effect models were used to correct for unwanted sources of variation, while attempting to preserve biological variation and accounting for potential heteroscedasticity. The pipeline was applied to targeted metabolomics data acquired in eight cancer-specific case-control studies nested within EPIC. Substantial inter-study and inter-batch heterogeneity was observed in the original data. Accordingly, the technical reproducibility was low-to-moderate for many metabolites with ICC values lower than 0.50, especially for the semiquantified metabolites (e.g., glycerophospholipids), suggesting that quantified metabolites might be less prone to unwanted variations due to analytical factors. Our normalization approach eliminated most of the inter-study and inter-batch variability, and improved the technical reproducibility of a large proportion of semi-quantified and quantified metabolites, with most ICC values greater than 0.75. Normalization using the ComBat approach^[19]^, which relies on a similar model but uses empirical Bayes estimation, performed similarly for all metabolites but acylcarnitines, for which ICC values were larger with our approach. ICC values estimated from the duplicate samples originating from different blood matrices (serum versus citrate plasma), were generally larger than 0.75 after normalization. However, they were also generally lower than values estimated from all duplicate samples. In particular, the ICC for methionine was 0.39 (95% confidence interval, CI: 0.14-0.57), as compared to 0.71 (95% CI: 0.64-0.77) when ICC estimation used all duplicate samples. This result calls for caution when pooling samples originating from different blood matrices, as large differences were reported for specific metabolites’ concentrations in serum and plasma samples^[26]^.

As samples within each individual EPIC study were all assayed in the same laboratory with the same LC-MS instruments, and mostly originated from the same blood matrix (except for GLBD that included serum and heparin plasma samples and BREA that included EDTA and citrate plasma samples), the variability due to these factors was encompassed into the inter-study variability, and could not be assessed by the PC-PR2 analysis. In particular, although the large inter-study variability in the non-normalized data supported the presence of inter-laboratory and inter-instrument variability, as previously reported for the AbsoluteIDQ p180 kit^[17]^, correction for batch and study effects also corrected for effects due to blood matrix and LC-MS instruments, which were both observed to contribute to less than 0.1% of the total variation in the normalized data. But, the inter-study and inter-batch variability also reflected biological variability, because factors like study center, gender, BMI and alcohol intake were not equally distributed across studies and batches. Consequently, some of the biological variation due to these factors would be removed if the normalized data were simply computed as the residuals in linear mixed models adjusted for study and batch. Conversely, by accounting for study center, gender, BMI and alcohol intake in the mixed models and by computing the normalized residuals using the step described in expression (2) in Section 4.2.4, the normalization preserved (some of) the variation due to these factors. This was illustrated by the distribution of normalized data that was shifted in opposite directions for studies including only men or women, and by the stronger associations with BMI observed when using the complete model for normalization compared to the simpler version that only included batch and study as random effects.

A critical step of normalization procedures that use linear mixed models, or more generally models with location/scale adjustments^[19]^, is the choice of (i) factors that may generate unwanted variation, for which a correction should be implemented, and (ii) factors that represent biological variability, which should be preserved after normalization. As illustrated in Section 4.2.4, while the list of variables in (i) should be included in matrix X (like study and batch), variables in (ii) should be included in matrix Z, and the choice depends on the study design and on the ultimate objective of the analysis. If the objective is to identify metabolites associated with a given phenotype, e.g., BMI, it is crucial to include BMI in matrix Z, particularly if BMI is associated with specific variables included in matrix X. Conversely, if the ultimate objective of the study is to identify metabolites associated with, say, alcohol, while controlling for BMI, then alcohol should be included in matrix Z (particularly if it is associated with specific variables included in matrix X), but BMI could be included in matrix X, so that the associations are adjusted for BMI. In any case, performing sensitivity analyses with normalized data generated including different sets of variables in matrices X and Z is a good practice.

In multicenter investigations like EPIC, study center is a sensitive variable, as it expresses technical (preanalytical) variation, likely the result of specific procedures for blood collection, sample treatment and storage, as well as biological variation reflecting specific lifestyle exposures, often characterized by geographical gradients. In addition, in multicenter context the relationship between two sets of variables could be evaluated at the overall level, at the center level or at the individual level^[27]^. In this study, to use the whole variability in metabolomics and BMI data, center was initially included in matrix Z. In sensitivity analysis, study center was included in matrix X, and the center-specific variability was removed. As shown in Supplementary Figure 4, results were similar to the overall analysis suggesting that group-level correlations were similar to individual–level correlations^[27]^. Alternative methods, like SVA^[28,20]^ and RRmix^[29]^, use linear (mixed) models with latent variables to estimate variability attributed to unspecified sources of variation, ultimately to be removed. These methods do not require prior knowledge of the sources of unwanted variation, but require the identification of sources of biological variation, as their effects would likely be removed if not properly accounted for in the linear predictor of the model.

The decision to implement data normalization largely depends on the ultimate objectives of the analysis. As the relationship between metabolites and cancer risk is generally quantified in conditional logistic regression models for matched case-control studies, metabolite measurements are compared within each matched case-control pair. If cases and controls are assayed within the same batch (as was the case in the EPIC metabolomics data), the effects of study and batch on the means of the measurements are not a concern, and normalization is not required, unless the variances of the measurements also vary across studies or batches. However, if the evaluation focuses on the investigation of lifestyle determinants of metabolomics data, like for example in mediation analysis, the matching is “broken” and control for inter-batch and inter-study variability is required^[7]^.

Although originally developed for the normalization of metabolomics data acquired in different studies, our pipeline could be used for data acquired in a single study, for example to correct for interbatch variability while preserving biological variability, and to correct for potential heteroscedastic structures of concentration levels across batches. Our pipeline could also be adapted to the normalization of biomarker data and other molecular data, possibly with some modifications. In particular, for the normalization of untargeted LC-MS metabolomics data, a step to exclude features based on comparison with blank samples should be added to the data cleaning ^[16]^, and a *K*-nearest neighbors approach has been shown to perform particularly well for the imputation of missing data^[15,30]^in the context of untargeted metabolomics data. Importantly, when processing untargeted metabolomics data from individual studies separately, different feature identifiers (e.g., mass to charge ratio and retention time) would characterize the same molecule in each study. Therefore, the pooling of several untargeted datasets would generally require an additional feature alignment step consisting in identifying the features present in the different datasets, which might be particularly challenging with data acquired in different laboratories^[31]^.

With the increasing availability of metabolomics data in large scale epidemiological investigations, such as those participating in the COnsortium of METabolomics Studies (COMETS)^[32]^, pooling will be more and more relevant as a strategy for increasing the statistical power when investigating the relationship between metabolomics data with disease indicators, environmental exposures and/or other -omics and biomarker data. Combined with analytical and graphical inspection of the data to determine sources of unwanted variability to be removed, and sources of biological variability to be preserved, linear mixed models provide a flexible tool to normalize metabolomics data, and possibly other -omics and biomarker data, prior to pooling data from different studies. As the comparability of measurements across studies is improved, our normalization approach could also be useful for studies that aims at the meta-analysis of individual-patient data from different studies, in particular if heteroscedastic patterns of variability were observed.

## 4. Materials and Methods

### 4.1. The EPIC study

EPIC is a large prospective study of over 500,000 men and women recruited in 1992-2000 in 23 centres in 10 European countries^*[21]*^, originally designed to investigate the relationship between diet and cancer risk. Incident cancer cases were identified through a combination of methods including linkage to health insurance records, cancer and pathology registries and active follow-up through study participants and their next-of-kin^*[21]*^. Around 386,000 participants from all countries provided a blood sample at recruitment. Fasting before blood withdrawal was not required. Blood was collected according to a standardized protocol in France, Germany, Greece, Italy, the Netherlands, Norway, Spain, and the UK^*[21]*^. Serum (except in Norway), plasma, erythrocytes, and buffy coat aliquots were stored in liquid nitrogen (−196 °C) in a centralized biobank at IARC. In Denmark, blood fractions were stored locally in the vapor phase of liquid nitrogen containers (−150 °C), and in Sweden, they were stored locally at −80 °C in standard freezers. Our analyses used targeted metabolomics data collected within the EPIC study and generated through the AbsoluteIDQ p180 or p150 commercial kit (Biocrates Life Science AG, Innsbruck Austria).

All participants provided written informed consent to participate in the EPIC study. This study was approved by the ethics committee of the International Agency for Research on Cancer (IARC) and all centers.

### 4.2. The pipeline to normalize data

Given a matrix of *p* metabolites acquired on *n* samples, our pipeline implemented four main steps, as summarized in Figure 7 and detailed hereafter for the EPIC targeted metabolomics data. R scripts implementing these four steps will be made available from GitHub.

**Figure 7.**
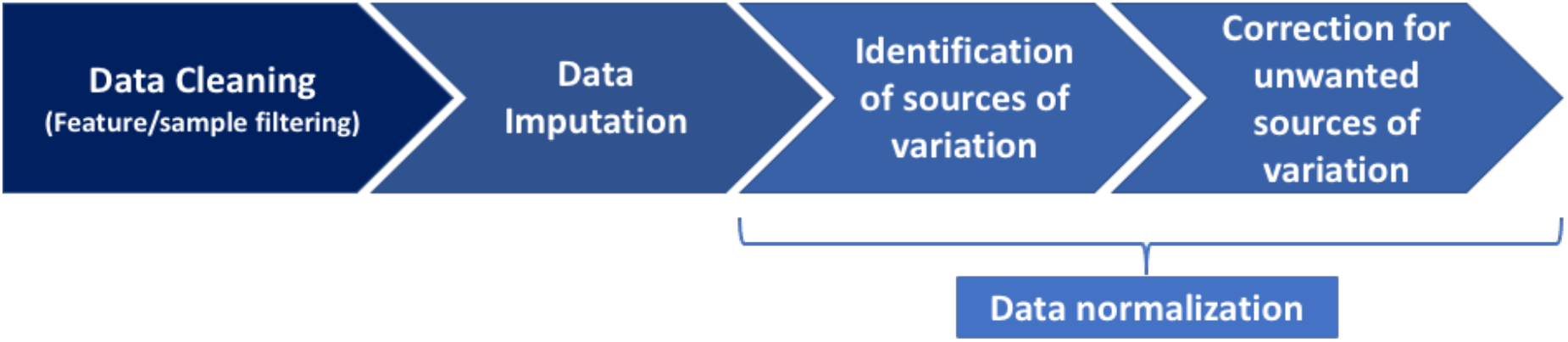
Main steps of the pipeline.

#### 4.2.1 Step1: Data cleaning

The objective of data cleaning was to remove the least informative metabolites and samples, using a number of (subjective) criteria. First, the pipeline excluded metabolites and samples exceeding a certain threshold of missingness (e.g., 20%), in each study separately. Missing values were here defined as fully missing values, for which no information on the real value was available. In particular they did not include out of measurable range values, which corresponded to values that were missing because they were below the batch-specific limit of detection (LOD), below the kit-specific lower limit of quantification (LLOQ), or above the kit-specific upper limit of quantification (ULOQ). An extra step was implemented to exclude outlying samples within each batch based on Principal Component Analysis (PCA)^[11]^, using a 20% proportional expansion of the Hotellings T2 distribution ellipse, with the level of the ellipse set to 100*(1-0.05)/N_b_ % and N_b_ the total number of batches. Samples assayed in batches with less than 10 samples were also excluded to ensure enough information during batch-specific data imputation (Section 4.2.2) and normalization (Section 4.2.4).

#### 4.2.2 Step 2: Data imputation

All missing values, including the out of measurable range values, were imputed in the cleaned dataset in each batch separately. Values below batch-specific LOD, below kit-specific LLOQ, or above kit-specific ULOQ were set to LOD/2, LLOQ/2 and ULOQ, respectively. Values below an unknown batch-specific LOD were set to LOD/2 after setting batch-specific LOD to study-specific medians of known LOD values. Fully missing values were set to the batch-specific median of non-missing values if less than 50% of the measurements in the batch were missing, and to the study-specific median of the batch-specific medians otherwise. Measurements were log-transformed to reduce skewness.

#### 4.2.3 Step 3: Data normalization, part 1: Identification of sources of variation

The PC-PR2 technique was used to identify main sources of variation in the metabolomics data^[13]^. The PC-PR2 is a multivariate technique that combines PCA with multiple linear regression to assess the proportion of the variability of the full metabolomics dataset explained by a set of explanatory variables, including samples characteristics (age, sex, BMI, alcohol consumption, study center), as well as preanalytical and analytical factors (fasting status, sample processing protocol, blood matrix, study, batch, laboratory instrument). While the former set of factors likely determined biological variability, the latter set likely introduced sources of unwanted variation in metabolomics data. PCA was conducted on metabolite measurements, and a number K≥1 of components sufficient to explain more than 80% of total variability was retained. Component scores were, in turn, regressed on the list of aforementioned independent variables, say W_1_, …,W_Q_, in multiple linear regression models, and the partial R^2^ for each covariate W_q_ was estimated for each component (C_k_). For example, the partial-R^2^ for W_1_ conditional on the (Q-1) other covariates for component k was

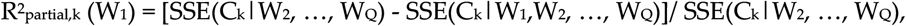

with SSE(C_k_|W_j_, …, W_Q_) expressing the residual sum of squares in the linear regression model of component C_k_ on variables W_j_, …, W_Q_, for j=1, 2. For variables with a nested structure, for example study (S) and batch within study (B), the formula was

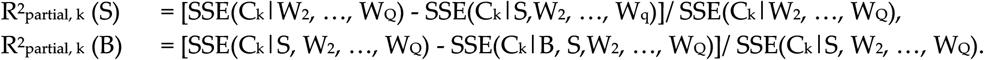

An overall R^2^_partial_ (W_1_) was obtained by the average of terms R^2^_partial,k_ (W_1_) weighted by the eigenvalue of each component. This overall estimate provides a measure of the variability in the ensemble of metabolite concentrations that each explanatory variable contributes to explain. The PC-PR2 technique is implemented in the pcpr2 R package available on GitHub.

#### 4.2.4 Step 4: Data normalization, part 2: Correction for the unwanted sources of variation

In order to correct for unwanted sources of variability while preserving biological variability, a random effects model was used for each metabolite separately^[14]^, as

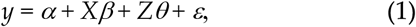

where *y* is the n-vector of the measurements for the metabolite/feature under consideration (all studies combined), the matrix *X* expresses variables corresponding to sources of variations that should be corrected for, and the optional matrix *Z* expresses variables corresponding to biological variations that should be preserved. Variables expressed in matrices X and Z typically include some of the variables W_1_, …, W_Q_ of the PC-PR2 analysis with largest R^2^_partial_. The vector of parameters *β* associated to matrix *X* may include both fixed- and random-effects, while the vector *θ* associated to matrix *Z* contains fixed effects only. Parameter *α* is the intercept, and vector *ε* ~ *N*_n_*(0, Σ)* corresponds to the random error of the model. Residuals *ε* are independent of the random effects of the model. Random effects are Gaussian, centered, and were further assumed to have diagonal covariance matrix in our illustration.

Parameters *α, β, θ* and the vector of residuals *ε* under model (1) are estimated by, say, *a*, *b, c*, and *e*. Normalized residual measurements are computed as

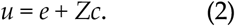

In this way, the normalization preserves the association between the metabolite and variables in *Z*, while any association with variables in *X* is eliminated. As mentioned in the Discussion, variables describing biological variations of interest should be included in matrix Z if they are associated with variables included in matrix X (e.g., sources of biological variations that are unequally balanced across studies or batches), otherwise some of the variation they explain would be removed because of the adjustment for X. In our illustration, study center indicators, gender, body mass index and alcohol intake were included in matrix Z, while batch and study indicators were included in matrix X.

In the simple homoscedastic random effect models, each component of the vector *ε* of residuals has the same variance: *Σ* = σ^2^*I_n_* for some σ^2^>0, where we denote by *I_p_* the identity matrix of size *p* for any positive integer *p*. However, in practice, pre-analytical and/or analytical factors may not only influence the means of the measurements via the term X *β* in model (1), but also their variance. For example, variances of components of *ε* may vary across studies. This was accounted for by working under heteroscedastic random effect models with a specified structure for the variance matrix *Σ* of the residuals, e.g., *Σ* was made of blocks of the form σ_*s*_^2^*I_ns_* for observations corresponding to study s (with *ns* the number of observations in study s). Then, residuals *e* were replaced by the Pearson residuals in Equation (2), after rescaling them to ensure that their overall variance equals that of the standard residuals. Homoscedastic models were implemented with the lmer function of the lme4 R package, while heteroscedastic models were implemented with the lme function of the nlme R package, using the *weights* instruction to specify the within-group heteroscedasticity structure.

For comparison, we also considered the ComBat method^[19]^ of the sva R package^[20]^, under which a fixed-effects version of model (1) is estimated using an empirical Bayes approach, to leverage the fact that sources of variation may affect many metabolites in similar ways. In our illustration, ComBat was applied to correct for batch effect (which also accounts for study effect), while attempting to preserve variations due to study center, gender, body mass index and alcohol intake.

### 4.3. Computation of the intra-class correlation coefficient using duplicated samples

The EPIC data included duplicate samples, corresponding to aliquots of a baseline blood sample from the same subject measured twice in different batches or in different studies. These duplicated samples were used to assess the technical reproducibility of metabolomics measurements, and in particular to compare technical reproducibility before and after normalization. In sensitivity analyses, ICC values were estimated using only duplicate samples originating from distinct blood matrices (serum and citrate plasma). For each metabolite, we estimated its ICC using a linear mixed effects model of the form^[16]^

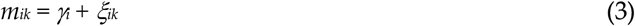

where *m_ik_* is the *k*-th replicate measurement of subject *i*, *k=1, 2, γ_i_ ~ N(μ, σγ^2^)* is a subject-specific random effect (with *μ* corresponding to the general mean of *m_ik_*), *ξ_ik_ ~ N(0, σξ^2^)* is the residual random error for replicate *k* of subject *i*, and Cov(*ξ_ik_*, *γ_i_*)=0. Under this model, ICC = Var(*γ_I_*)/Var(*γ_I_ + ξ_ik_*), so the ICC estimate was defined as the ratio of the estimated between-subject variance to the estimated total variance (between- and within-subject). Model (3) above can be estimated even if only a portion of the subjects have replicated samples. It was implemented using the lmer function of the lme4 R package, and 95% confidence intervals (CI) of the ICC values were derived using the parametric bootstrap implemented by the bootMer function of the lme4 R package.

## Author Contributions

Conceptualization, V.V., M.H. and P.F.; methodology V.V. and P.F.; software, V.V.; validation, V.V., M.H., S.R.; resources, B.H., M.Jo., L.D., M.G., M.Je., S.R., R.T., J.S., L.B.; data curation, B.H., V.V.; writing–original draft preparation, V.V., M.H., S.R., M.B., L.D., R.S., P.R.K., P.F.; writing–review and editing, all co-authors; All authors have read and agreed to the published version of the manuscript.

## Funding

The coordination of EPIC is financially supported by International Agency for Research on Cancer (IARC) and by the Department of Epidemiology and Biostatistics, School of Public Health, Imperial College London which has additional infrastructure support provided by the NIHR Imperial Biomedical Research Centre (BRC). The national cohorts are supported by: Danish Cancer Society (Denmark); Ligue Contre le Cancer, Institut Gustave Roussy, Mutuelle Générale de l’Education Nationale, Institut National de la Santé et de la Recherche Médicale (INSERM) (France); German Cancer Aid, German Cancer Research Center (DKFZ), German Institute of Human Nutrition Potsdam-Rehbruecke (DIfE), Federal Ministry of Education and Research (BMBF) (Germany); Associazione Italiana per la Ricerca sul Cancro-AIRC-Italy, Compagnia di SanPaolo and National Research Council (Italy); Dutch Ministry of Public Health, Welfare and Sports (VWS), Netherlands Cancer Registry (NKR), LK Research Funds, Dutch Prevention Funds, Dutch ZON (Zorg Onderzoek Nederland), World Cancer Research Fund (WCRF), Statistics Netherlands (The Netherlands); Health Research Fund (FIS) - Instituto de Salud Carlos III (ISCIII), Regional Governments of Andalucía, Asturias, Basque Country, Murcia and Navarra, and the Catalan Institute of Oncology - ICO (Spain); Swedish Cancer Society, Swedish Research Council and County Councils of Skåne and Västerbotten (Sweden); Cancer Research UK (14136 to EPIC-Norfolk; C8221/A29017 to EPIC-Oxford), Medical Research Council (1000143 to EPIC-Norfolk; MR/M012190/1 to EPIC-Oxford) (United Kingdom). IDIBELL acknowledges support from the Generalitat de Catalunya through the CERCA Program. R.Z.-R. would like to thank the “Miguel Servet” program (CPII20/00009) from the Institute of Health Carlos III (Spain) and the European Social Fund (ESF).The breast cancer study (BREA) was funded by the French National Cancer Institute (grant number 2015-166). The colorectal cancer studies (CLRT1 and CRLT2) were funded by World Cancer Research Fund (MG; reference: 2013/1002; www.wcrf.org/), the European Commission (MG; FP7: BBMRI-LPC; reference: 313010; https://ec.europa.eu/). The endometrial cancer study (ENDO) was funded by Cancer Research UK (grant number C19335/A21351). The kidney study (KIDN) was funded by the World Cancer Research Fund (MJ; reference: 2014/1193; www.wcrf.org/) and the European Commission (MJ; FP7: BBMRI-LPC; reference: 313010; https://ec.europa.eu/). The generation of metabolomics data in the gallbladder cancer study (GLBD) was supported by the European Union within the initiative “Biobanking and Biomolecular Research Infrastructure—Large Prospective Cohorts” (Collaborative study “Identification of biomarkers for gallbladder cancer risk prediction—Towards personalized prevention of an orphan disease”) under grant agreement no. 313010 (BBMRI-LPC). The liver cancer study (LIVE) was supported in part by the French National Cancer Institute (L’Institut National du Cancer; INCa; grant numbers 2009-139 and 2014-1-RT-02-CIRC-1; PI: M. Jenab) and by internal funds of the IARC. For the participants in the prostate cancer study (PROS), sample retrieval and preparation, and assays of metabolites were supported by Cancer Research UK (C8221/A19170), and funding for grant 2014/1183 was obtained from the World Cancer Research Fund (WCRF UK), as part of the World Cancer Research Fund International grant programme. Mathilde His’ work reported here was undertaken during the tenure of a postdoctoral fellowship awarded by the International Agency for Research on Cancer, financed by the Fondation ARC.

## Conflicts of Interest

The authors declare no conflict of interest.

## IARC disclaimer

Where authors are identified as personnel of the International Agency for Research on Cancer/World Health Organization, the authors alone are responsible for the views expressed in this article and they do not necessarily represent the decisions, policy, or views of the International Agency for Research on Cancer/World Health Organization.

---

**Supplementary Figure 1.**
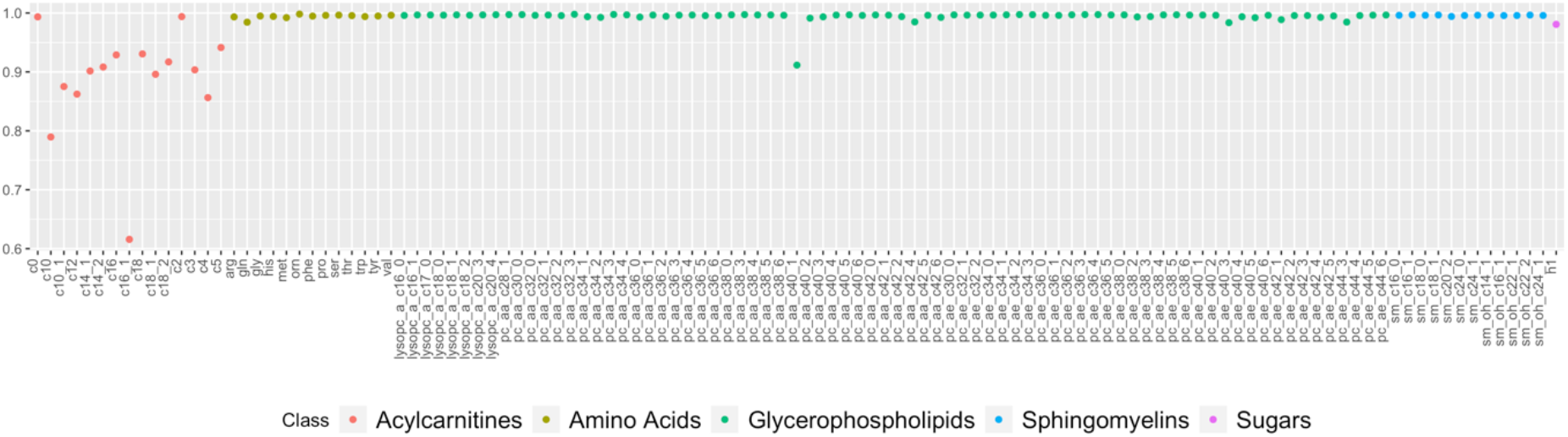
Correlations between normalized measurements produced by ComBat and our approach. Both approaches were run to correct for batch and study effects, and to preserve biological variations due to study center, gender, alcohol intake and body mass index.

**Supplementary Figure 2.**
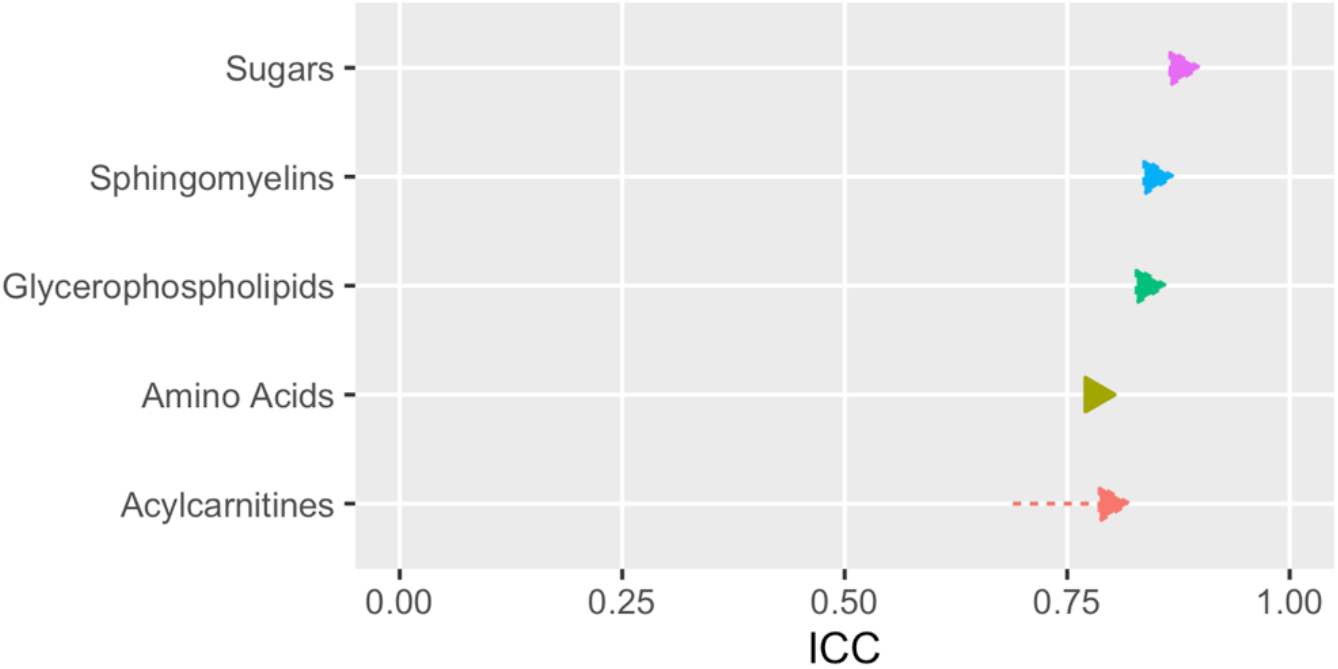
Average ICC values for each class of metabolites after normalization using ComBat and our approach; for each arrow, its origin represents the ICC obtained when using ComBat for normalization and its peak represents the ICC obtained when using our approach for normalization. All arrows are oriented towards the right, especially for acylcarnitins, indicating that our approach produced more reproducible measurements for most metabolites.

**Supplementary Figure 3.**
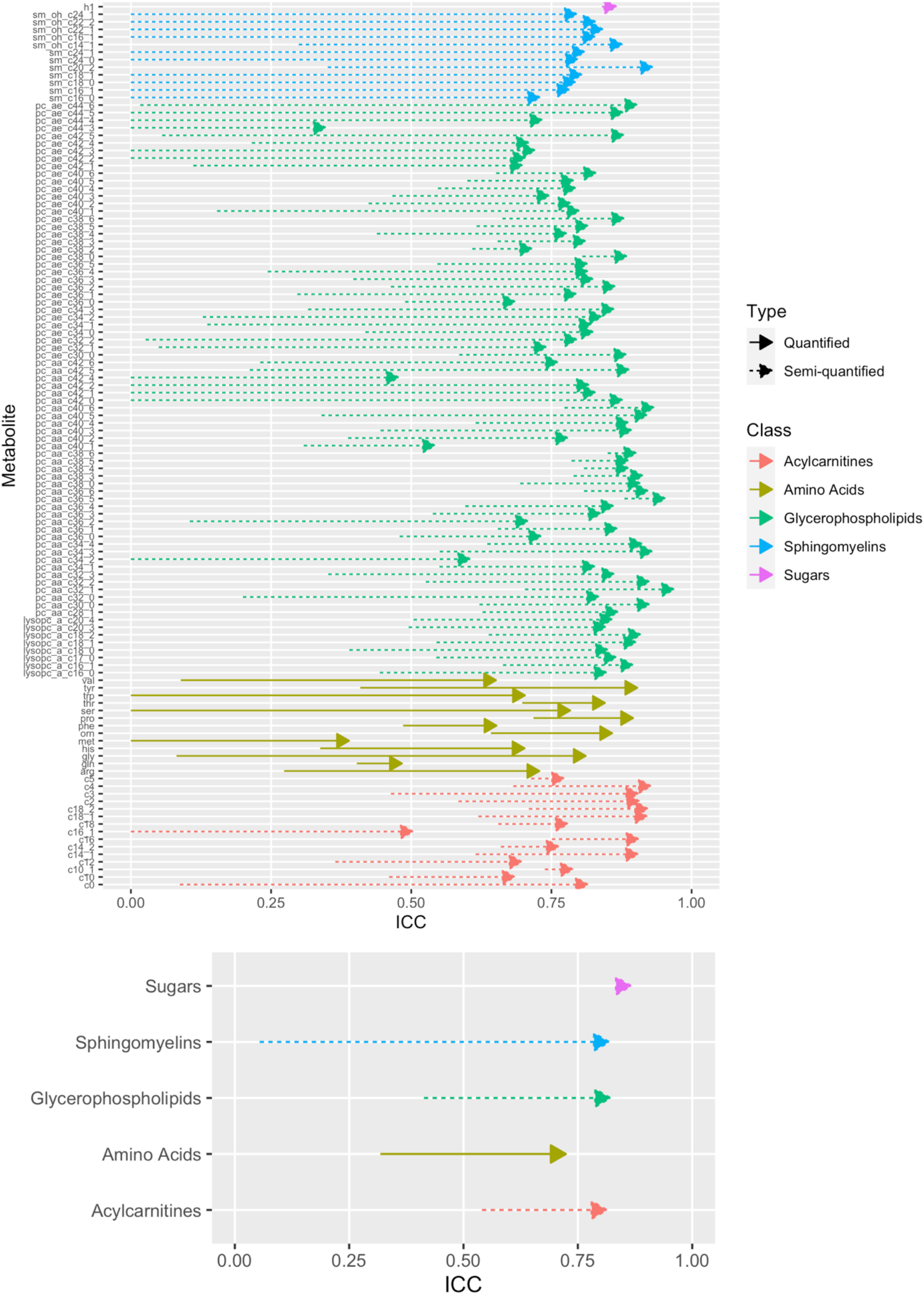
Metabolite-specific ICC values before and after normalization (top) and average ICC values for each class of metabolites before and after normalization (bottom); normalization was conducted so as to remove study and batch effects while preserving variation due to study center, BMI, gender and alcohol intake. Only duplicate samples measured in two different studies and originating from two different blood matrices (serum and citrate plasma) were used here. For each arrow, its origin represents the ICC value before normalization, and its peak represents the ICC value after normalization.

**Supplementary Figure 4.**
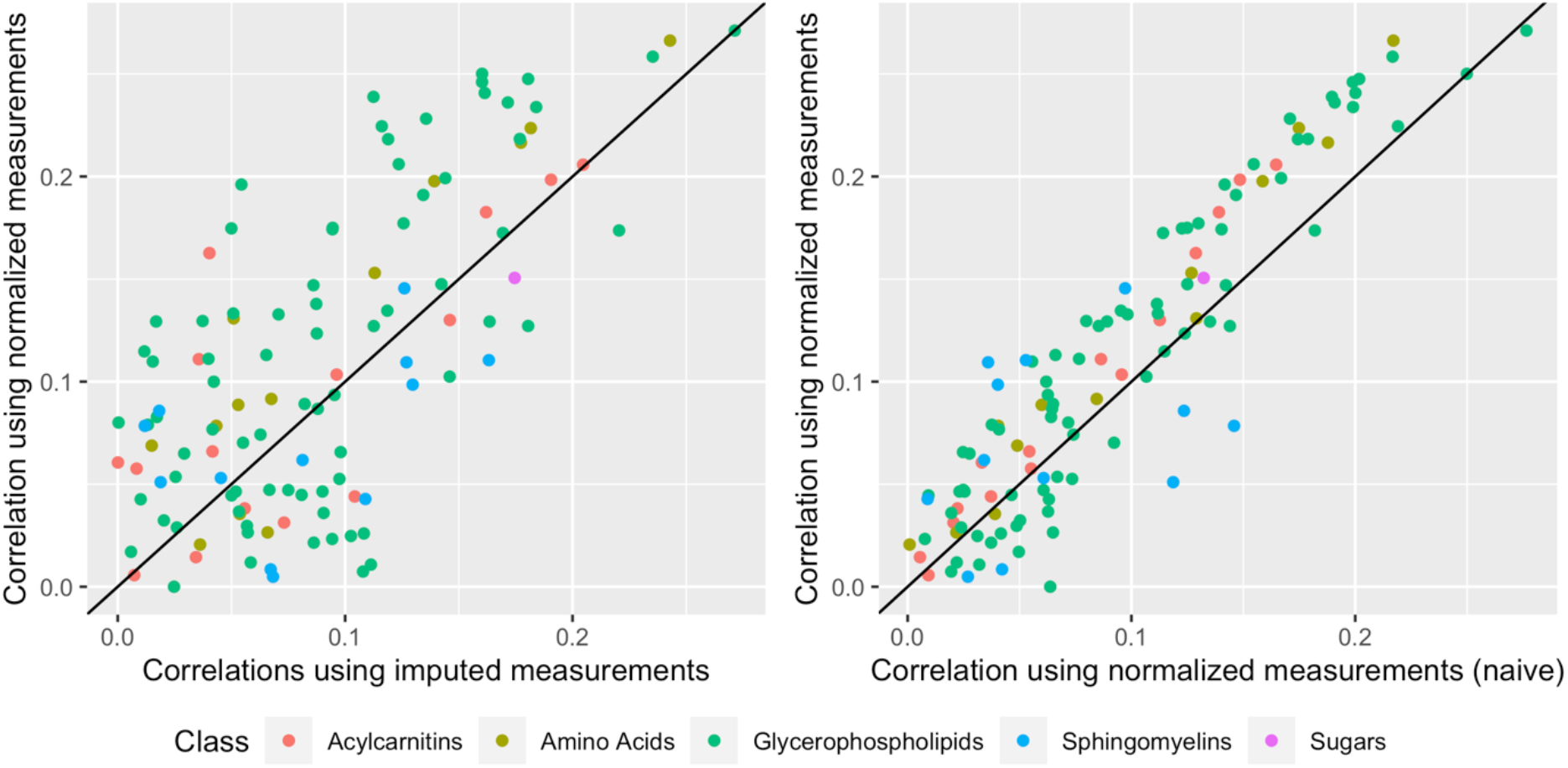
Correlations (absolute values) between BMI and the 117 metabolites in control samples. The *y*-axis represents values computed with normalized measurements (the normalization was run so as to remove study, batch and center effects while preserving variation due to BMI, gender and alcohol intake), while the *x*-axis represents values computed with imputed (non-normalized) measurements (left), and normalized measurements produced by the “naïve” normalization (right), which corrects for study, batch and center effects without preserving variation due to BMI, gender and alcohol intake.

**Supplementary Table 1.** List of the 117 metabolites retained after the data cleaning step.

| Name | Symbol in Figures | Class |
| --- | --- | --- |
| Carnitine | c0 | Acylcarnitins |
| Acetylcarnitine | c2 | Acylcarnitins |
| Propionylcarnitine | c3 | Acylcarnitins |
| Butyrylcarnitine | c4 | Acylcarnitins |
| Valerylcarnitine | c5 | Acylcarnitins |
| Decanoylcarnitine | c10 | Acylcarnitins |
| Decenoylcarnitine | c10_1 | Acylcarnitins |
| Dodecanoylcarnitine | c12 | Acylcarnitins |
| Tetradecenoylcarnitine | c14_1 | Acylcarnitins |
| Tetradecadienylcarnitine | c14_2 | Acylcarnitins |
| Hexadecanoylcarnitine | c16 | Acylcarnitins |
| Hexadecenoylcarnitine | c16_1 | Acylcarnitins |
| Octadecanoylcarnitine | c18 | Acylcarnitins |
| Octadecenoylcarnitine | c18_1 | Acylcarnitins |
| Octadecadienylcarnitine | c18_2 | Acylcarnitins |
| Arginine | arg | Amino Acids |
| Glutamine | gln | Amino Acids |
| Glycine | gly | Amino Acids |
| Histidine | his | Amino Acids |
| Methionine | met | Amino Acids |
| Ornithine | orn | Amino Acids |
| Phenylalanine | phe | Amino Acids |
| Proline | pro | Amino Acids |
| Serine | ser | Amino Acids |
| Threonine | thr | Amino Acids |
| Tryptophan | trp | Amino Acids |
| Tyrosine | tyr | Amino Acids |
| Valine | val | Amino Acids |
| lysoPC a C16:0 | lysopc_a_c16_0 | Glycerophospholipids |
| lysoPC a C16:1 | lysopc_a_c16_1 | Glycerophospholipids |
| lysoPC a C17:0 | lysopc_a_c17_0 | Glycerophospholipids |
| lysoPC a C18:0 | lysopc_a_c18_0 | Glycerophospholipids |
| lysoPC a C18:1 | lysopc_a_c18_1 | Glycerophospholipids |
| lysoPC a C18:2 | lysopc_a_c18_2 | Glycerophospholipids |
| lysoPC a C20:3 | lysopc_a_c20_3 | Glycerophospholipids |
| lysoPC a C20:4 | lysopc_a_c20_4 | Glycerophospholipids |
| PC aa C28:1 | pc_aa_c28_1 | Glycerophospholipids |
| PC aa C30:0 | pc_aa_c30_0 | Glycerophospholipids |
| PC aa C32:0 | pc_aa_c32_0 | Glycerophospholipids |

**Supplementary Table 1 (continued)**
| Name | Symbol in Figures | Class |
| --- | --- | --- |
| PC aa C32:1 | pc_aa_c32_1 | Glycerophospholipids |
| PC aa C32:3 | pc_aa_c32_3 | Glycerophospholipids |
| PC aa C34:1 | pc_aa_c34_1 | Glycerophospholipids |
| PC aa C34:2 | pc_aa_c34_2 | Glycerophospholipids |
| PC aa C34:3 | pc_aa_c34_3 | Glycerophospholipids |
| PC aa C34:4 | pc_aa_c34_4 | Glycerophospholipids |
| PC aa C36:0 | pc_aa_c36_0 | Glycerophospholipids |
| PC aa C36:1 | pc_aa_c36_1 | Glycerophospholipids |
| PC aa C36:2 | pc_aa_c36_2 | Glycerophospholipids |
| PC aa C36:3 | pc_aa_c36_3 | Glycerophospholipids |
| PC aa C36:4 | pc_aa_c36_4 | Glycerophospholipids |
| PC aa C36:5 | pc_aa_c36_5 | Glycerophospholipids |
| PC aa C36:6 | pc_aa_c36_6 | Glycerophospholipids |
| PC aa C38:0 | pc_aa_c38_0 | Glycerophospholipids |
| PC aa C38:3 | pc_aa_c38_3 | Glycerophospholipids |
| PC aa C38:4 | pc_aa_c38_4 | Glycerophospholipids |
| PC aa C38:5 | pc_aa_c38_5 | Glycerophospholipids |
| PC aa C38:6 | pc_aa_c38_6 | Glycerophospholipids |
| PC aa C40:1 | pc_aa_c40_1 | Glycerophospholipids |
| PC aa C40:2 | pc_aa_c40_2 | Glycerophospholipids |
| PC aa C40:3 | pc_aa_c40_3 | Glycerophospholipids |
| PC aa C40:4 | pc_aa_c40_4 | Glycerophospholipids |
| PC aa C40:5 | pc_aa_c40_5 | Glycerophospholipids |
| PC aa C40:6 | pc_aa_c40_6 | Glycerophospholipids |
| PC aa c42:0 | pc_aa_c42_0 | Glycerophospholipids |
| PC aa c42:1 | pc_aa_c42_1 | Glycerophospholipids |
| PC aa C42:2 | pc_aa_c42_2 | Glycerophospholipids |
| PC aa C42:4 | pc_aa_c42_4 | Glycerophospholipids |
| PC aa C42:5 | pc_aa_c42_5 | Glycerophospholipids |
| PC aa C42:6 | pc_aa_c42_6 | Glycerophospholipids |
| PC ae C30:0 | pc_ae_c30_0 | Glycerophospholipids |
| PC ae C30:1 | pc_ae_c32_1 | Glycerophospholipids |
| PC ae C30:2 | pc_ae_c32_2 | Glycerophospholipids |
| PC ae C34:0 | pc_ae_c34_0 | Glycerophospholipids |
| PC ae C34:1 | pc_ae_c34_1 | Glycerophospholipids |
| PC ae C34:2 | pc_ae_c34_2 | Glycerophospholipids |
| PC ae C34:3 | pc_ae_c34_3 | Glycerophospholipids |
| PC ae C36:0 | pc_ae_c36_0 | Glycerophospholipids |

**Supplementary Table 1 (continued)**
| Name | Symbol in Figures | Class |
| --- | --- | --- |
| PC ae C36:1 | pc_ae_c36_1 | Glycerophospholipids |
| PC ae C36:2 | pc_ae_c36_2 | Glycerophospholipids |
| PC ae C36:3 | pc_ae_c36_3 | Glycerophospholipids |
| PC ae C36:4 | pc_ae_c36_4 | Glycerophospholipids |
| PC ae C36:5 | pc_ae_c36_5 | Glycerophospholipids |
| PC ae C38:0 | pc_ae_c38_0 | Glycerophospholipids |
| PC ae C38:2 | pc_ae_c38_2 | Glycerophospholipids |
| PC ae C38:3 | pc_ae_c38_3 | Glycerophospholipids |
| PC ae C38:4 | pc_ae_c38_4 | Glycerophospholipids |
| PC ae C38:5 | pc_ae_c38_5 | Glycerophospholipids |
| PC ae C38:6 | pc_ae_c38_6 | Glycerophospholipids |
| PC ae C40:1 | pc_ae_c40_1 | Glycerophospholipids |
| PC ae C40:2 | pc_ae_c40_2 | Glycerophospholipids |
| PC ae C40:3 | pc_ae_c40_3 | Glycerophospholipids |
| PC ae C40:4 | pc_ae_c40_4 | Glycerophospholipids |
| PC ae C40:5 | pc_ae_c40_5 | Glycerophospholipids |
| PC ae C40:6 | pc_ae_c40_6 | Glycerophospholipids |
| PC ae C42:1 | pc_ae_c42_1 | Glycerophospholipids |
| PC ae C42:2 | pc_ae_c42_2 | Glycerophospholipids |
| PC ae C42:3 | pc_ae_c42_3 | Glycerophospholipids |
| PC ae C42:4 | pc_ae_c42_4 | Glycerophospholipids |
| PC ae C42:5 | pc_ae_c42_5 | Glycerophospholipids |
| PC ae C44:3 | pc_ae_c44_3 | Glycerophospholipids |
| PC ae C44:4 | pc_ae_c44_4 | Glycerophospholipids |
| PC ae C44:5 | pc_ae_c44_5 | Glycerophospholipids |
| PC ae C44:6 | pc_ae_c44_6 | Glycerophospholipids |
| SM C16:0 | sm_c16_0 | Sphingomyelins |
| SM C16:1 | sm_c16_1 | Sphingomyelins |
| SM C18:0 | sm_c18_0 | Sphingomyelins |
| SM C18:1 | sm_c18_1 | Sphingomyelins |
| SM C20:2 | sm_c20_2 | Sphingomyelins |
| SM C24:0 | sm_c24_0 | Sphingomyelins |
| SM C24:1 | sm_c24_1 | Sphingomyelins |
| SM (OH) C14:1 | sm_oh_c14_1 | Sphingomyelins |
| SM (OH) C16:1 | sm_oh_c16_1 | Sphingomyelins |
| SM (OH) C22:1 | sm_oh_c22_1 | Sphingomyelins |
| SM (OH) C22:2 | sm_oh_c22_2 | Sphingomyelins |
| SM (OH)C24:1 | sm_oh_c24_1 | Sphingomyelins |
| Hexoses | h1 | Monosaccharides |

**Supplementary Table 2:** Study origin of duplicate samples in the EPIC targeted metabolomics data.

| Study 1 | Study 2 | Number of EPIC participants |
| --- | --- | --- |
| BREA | CLRT1 | 1 |
| BREA | CLRT2 | 4 |
| BREA | ENDO | 5 |
| BREA | GLBD | 1 |
| BREA | LIVE | 2 |
| CLRT1 | CLRT2 | 2 |
| CLRT1 | ENDO | 2 |
| CLRT1 | KIDN | 2 |
| CLRT1 | LIVE | 1 |
| CLRT1 | PROS | 4 |
| CLRT2 | ENDO | 4 |
| CLRT2 | KIDN | 5 |
| CLRT2 | PROS | 27 |
| ENDO | KIDN | 2 |
| GLBD | LIVE | 51 |
| GLBD | PROS | 1 |
| KIDN | LIVE | 1 |
| KIDN | PROS | 23 |
| LIVE | PROS | 9 |

## Notes

### Competing Interest Statement

The authors have declared no competing interest.

